# Gene expression asymmetry in Parkinson’s Disease; variation of *CCT* and *BEX* gene expression levels are correlated with hemisphere specific severity

**DOI:** 10.1101/2024.07.02.601704

**Authors:** Steven E. Pierce, Edwin J.C. van der Schans, Elizabeth Ensink, Gerhard A. Coetzee

## Abstract

Parkinson’s Disease (PD) develops unilaterally, which may be related to brain hemispheric differences in gene expression. Here we measured bulk RNA-seq levels in neuronal nuclei obtained from prefrontal cortex postmortem brain samples from males and females with PD and from healthy controls. Left and right hemispheres from each brain were related the side of symptom onset and compared. We employed two *a priori* approaches; first we identified genes differentially expressed between PD and controls and between left vs right PD brain hemispheres. Second, we examined the presence of, and correlates to, variable asymmetry seen in candidate PD differentially expressed genes. We found large variation among individuals with PD, and PD stratification by gene expression similarity was required for patterns of genetic asymmetry to emerge. For a subset of PD brains, hemispherical variation of *CCT* and *BEX* gene levels correlated with the side of PD symptom onset.

## Introduction

Asymmetric pathology is a well-documented aspect of many neurodegenerative diseases [1, 2]. Parkinson’s Disease (PD) presents clinically obvious and persistent right/left differences in both motor and nonmotor PD symptoms [1, 3]. Correspondingly, diagnostic differences between hemispheres are obvious and persistent including through dopamine transporter imaging or diffusion MRI [4, 5]. These phenomena correspond to dopaminergic neuronal loss and other pathological signatures in more affected (contralateral) brain hemispheres apparent in postmortem tissue [6].

PD asymmetry is not entirely random. For instance, there is a small but significant bias for onset side to correspond to the dominant hand [7, 8]. Any amount of non-randomness indicates some intrinsic or extrinsic brain hemisphere differences exists and are relevant to onset and/or progression of PD. To put it another way, given persistent nonrandom asymmetry exists, either one hemisphere is relatively protected (compared to the other) from early on, or one hemisphere becomes relatively protected (i.e. is less burdened) after disease onset. Identifying the underpinning of either or both mechanisms would be valuable for diagnostic and treatment strategies.

There are two possibilities for nonrandom PD asymmetry, which are not mutually exclusive [1]. First, intrinsic differences or systematic lateralized environmental exposures may cause one hemisphere to be more vulnerable to the disease than the other. The second possibility is that differences may be *induced by* the disease. Vulnerability to PD may be equivalent between hemispheres at some arbitrary developmental stage, but in the absence of a synchronizing mechanism, asymmetry could persist or become more exaggerated following onset. Stochastic processes likely lead to subtly different disease burden to either hemisphere. However, initially random effects could accumulate and potentially lead to amplifying feedback due to the dynamic interaction between localized and global disease responses. For example, the second type of process could involve a general protective stress response or neurogenic signal that is more effective or stronger in the less affected hemisphere, or a response with both positive and negative effects, such as inflammation, could be less beneficial and more pathological in the more effected hemisphere.

Recently, we and others from our institute lead by the late Dr. Viviane Labrie obtained and studied a set of post-mortem brains from control subjects and patients diagnosed with Parkinson’s Disease (PD). Useful for the study reported here, both hemispheres of the same brain were available including an indication of the side of PD symptom onset in the diseased cases. In the present study we compared bulk RNA-seq in neuronal nuclei from post-mortem brains categories PD, controls, male, female left-right hemispheres (related to PD side of symptom onset defined as moderate and severe hemisphere). The questions we sought to address were:

- Can gene expression differences be related to the brain hemispheric side of PD onset?
- Do the differences between moderate and severe hemispheres as unmatched and matched to symptoms, respectively, mimic healthy vs PD brains or are there novel differences?
- Are there any gene expression differences involved in brain lateralization or other early processes, that are likely to predispose one side to PD symptom onset?

## Methods

### Study Cohort

Frozen tissue samples were obtained through the NIH brain bank from the Human Brain and Spinal Fluid Resource Center in Los Angeles, CA and from the University of Miami Miller School of Medicine with approval from the ethics committees of the Van Andel Research Institute. For each individual, we had information on the donor’s age, sex, ethnicity, and cause of death. Additionally, the tissue post-mortem interval, duration of disease, side of onset, and neurofibrillary tangle burden (Braak stage[9]). Neurons of the prefrontal cortex were selected for this study, as this region shows pathology only at very advanced stages of PD and would be unlikely to show indicators of local neuronal loss. Matched hemisphere samples from 5 female and 5 male healthy control, and 9 female and 28 male PD brain samples were available for analysis. Braak stages ranged from 1.5 to 6 with a mean of 2.5. The average age at death was 76.1 years.

### RNA-sequencing

Neuronal nuclei were isolated from the human prefrontal cortex using flow cytometry and filtering for an anti-NeuN antibody. Neuronal nuclei mRNA provides a good representation of total mRNA[10]. Libraries were prepared by the Van Andel Genomics Core from 500 ng of total RNA using the KAPA RNA HyperPrep Kit with RiboseErase. RNA was sheared to an average of 300 bp. Individually indexed cDNA libraries were pooled, and 75-bp paired-end sequencing was performed on an Illumina NextSeq 500 sequencer. Base calling was done by Illumina NextSeq Control Software (NCS; v2.0), and the output of NCS was demultiplexed and converted to FastQ format with Illumina Bcl2fastq (v1.9.0)

### Differential Expression Analysis

Following sequencing, fastq files were trimmed using Trimmomatic V0.36 and the settings: ILLUMINACLIP:TruSeq3-PE.fa:2:30:10 LEADING:3 TRAILING:3 SLIDINGWINDOW:4:15 MINLEN:36. Trimmed fastq files were aligned to HG19 using STAR v2.7 [11]. RNA-seq: Alignments (bam files) were converted to feature counts using HTSeq v0.6.0 referenced against the ENSEMBLE annotation of HG19: Homo_sapiens.GRCh37.87.gtf counting against the feature “exon”, grouped by “gene_id”, and using the strand parameter “reverse”. Technical replicates were combined for samples with duplicates. Gene counts were normalized using edgeR (TMM) and tested for significant differential expression with Limma and Voom, in R (v4.10.2) [12-14]. Initial differential expression was estimated with a design of Gene expression = ∼0 + combined category (PD or hcntrl, sex, side of onset, R or L hemisphere) + duration of illness + library RIN + library size + difference relative to mean age + positive nuclei per mg tissue + PMI. This comparison was used to identify the least statistically different 2000 genes for healthy control vs. PD. The function RUVr via Ruvseq [15] was used to estimate 5 correcting variables to minimize residuals based on model: Gene expression = ∼0 + combined category + duration of illness + difference from mean age. These 5 normalizing variables were added to the second model for final analysis. For comparison of display of expression values the normalized count from RUVseq was used, though statistical analysis was based solely on logCPM from Limma. DE significance is defined by a FDR < 0.05 unless otherwise detailed. Data available under GSE264050.

### Enrichment Analysis

Enrichment of ontology for gene sets was done using ranked enrichment by FDR with GSEA [16], or for gene set enrichment by STRING V.12 [17].

## Results

To identify potential gene expression differences between the more affected (severe) and less affected (moderate) hemispheres, defined as matched and unmatched, respectively in PD brain tissue and in comparison, to healthy control samples, we performed bulk-RNA-sequencing on neuronal nuclei (NeuN+) derived from frontal cortex tissue of 28 PD brains and 10 healthy control brains. Paired samples from the left and right hemispheres of each brain were used. Paired hemisphere RNA-seq for 5 female and 5 male healthy control (average age 78, 69 years, respectively) were compared to data from 9 female (3 left onset, 5 right onset, 1 bilateral; ave. age, 77 years) and 28 male (9 left onset, 8 right onset, 2 bilateral; ave. age, 77 years) samples. The average Braak stage for PD brains was from stage 2-3.

We compared the gene expression levels in PD patients to that of healthy controls according to sex, both jointly and separately and including covariates for age, duration of illness, and 5 correcting variables. We found large expression changes (Fig1A). As expected, both sex specific and non-specific PD changes were identified. Additionally, the most prominent disease relevant processes based on gene set enrichment were driven by the expression of CCT chaperonin complex subunits (FigA,B; Supplemental Fig1A) The *CCT* genes express proteins that form the CCT chaperonin complex, which has been reported to be associated with Parkinson’s disease (PD) risk as well as being critical in PD processes including alpha-synuclein aggregation [18-20]. These genes, as well as other chaperone protein and protein folding related genes, were generally elevated in PD patients compared to healthy controls. Genes down-regulated in PD brains were less enriched for processes, but included axoneme and dynein linked genes, including *odad1-4* (Fig1C; Supplemental Fig1B).

**Figure 1.**
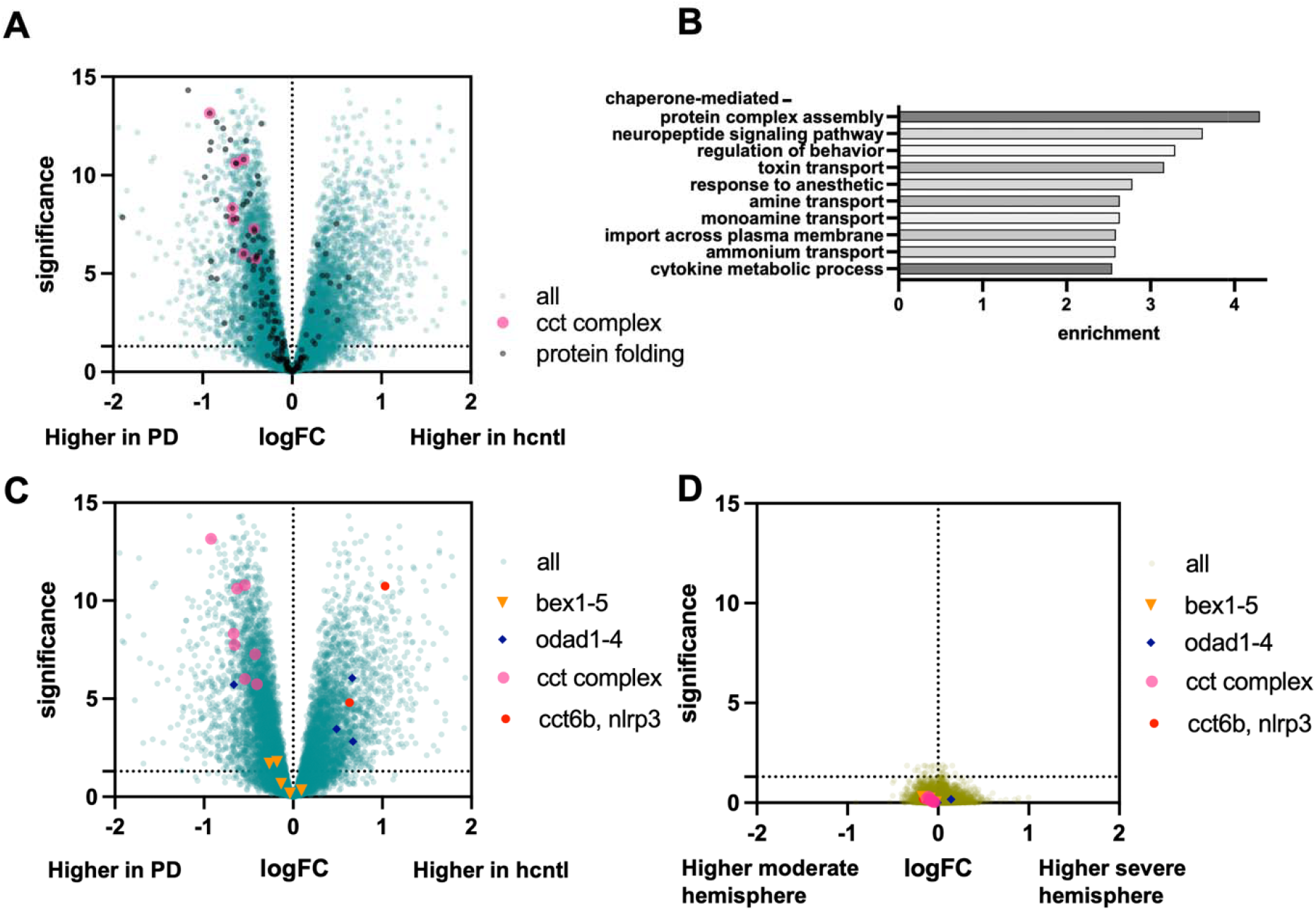
Differential gene expression across entire sample set. A) Volcano plot of DE comparing all PD samples vs. all healthy control samples (hcntl). All genes annotated with the biological process GO category “protein folding” are indicated in black, the 8 primary subunits of the CCT complex in magenta. B) the most highly enriched GO categories of all significant (FDR > 0.05) genes. C) Volcano as in A, but with 4 groups of genes indicated for use as marker posts and genes of interest to compare across different analysis. D) Volcano plot of DE comparing asymmetry (severe vs. moderate hemisphere) across all PD samples.

The central question of this study was what (if any) gene expression is statistically different across hemispheres relative to PD side of onset. In the first instance, we found little evidence that any genes were significantly asymmetrical by comparing severe to moderate hemispheres across all patients and by linear modeling including age, duration of illness, and 5 composite nuisance variables (Fig1d). A few genes were significant for males or females when analyzed separately (9 and 40 respectively, (Supplemental Fig1C).

Given this generally negative result, we sought further clarification to support or reject the initial observation that no asymmetrically expressed genes were present. The lack of straight forwardly significant asymmetry across the 25 (unilateral onset) PD brains could reflect the large variation between people compared to near identity across hemispheres within a given individual. Of course, a large enough study size may help alleviate this issue. As a more tractable tactic to further address variation between subjects, we asked whether any asymmetry was apparent after selecting only the most similar PD patients (based on gene expression levels) for comparison with controls. We clustered samples using rank correlation of the top 2500 most variable and highly expression genes. On this basis selected paired samples from the most similar cluster of PD only brains, consisting of 14 PD brains with both hemispheres (8 male, 6 female) were compared to the 10 control brain samples (Supplemental Fig2). In this narrower comparison, differentially expressed (DE) genes for PD vs. healthy control remained similar to the full comparison. However, unlike in the full PD group among a subset of similar PD brains, we identified many significant DE asymmetry genes comparing severe vs moderate hemispheres (Fig2C,D).

**Figure 2.**
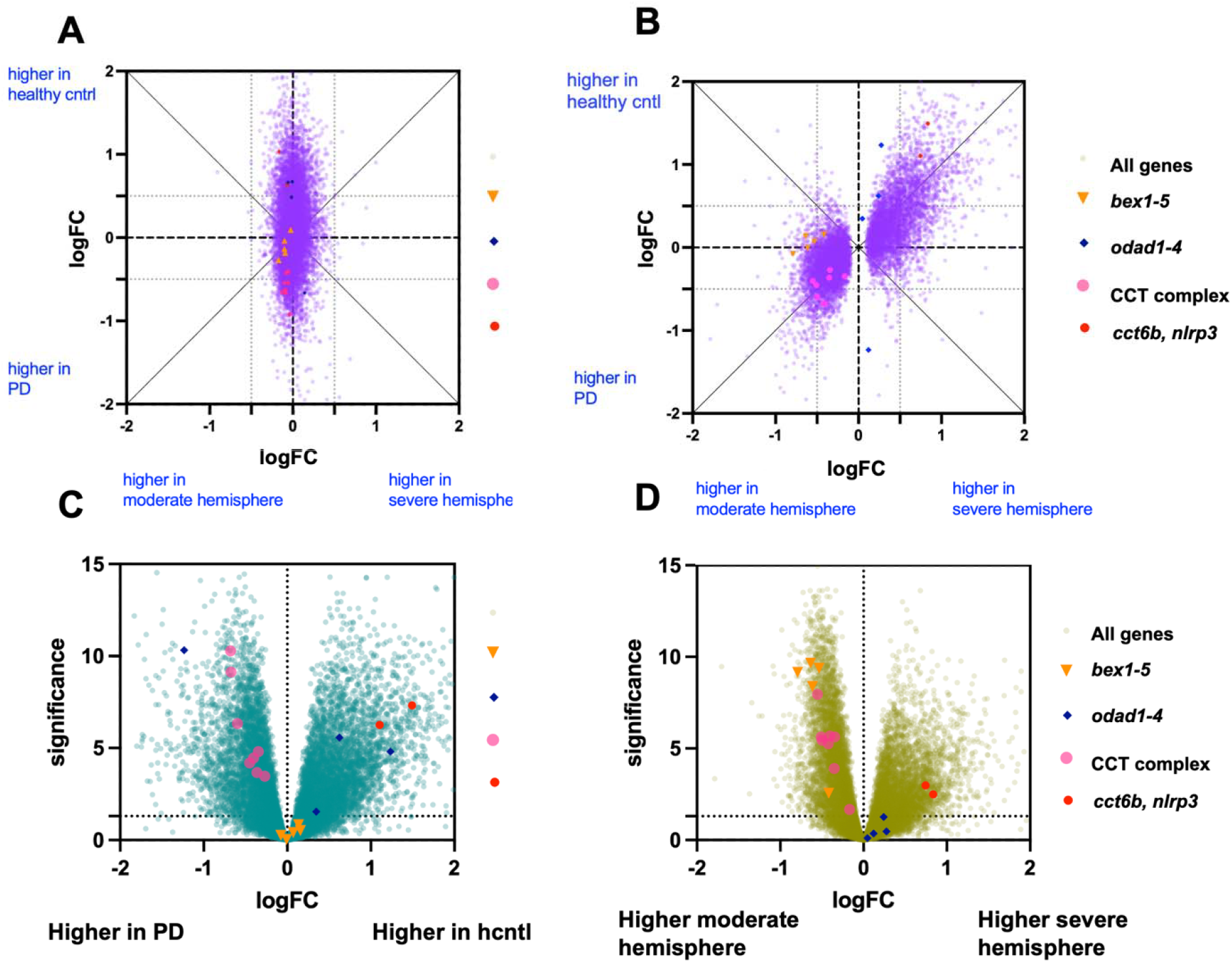
Differential gene expression across a subset of the most highly correlated PD brains. In purple are comparison of DE changes by disease state vs. asymmetry. GLM estimated log2 fold change of two separate comparisons: healthy control vs. PD (Y-axis) against severe hemisphere vs. moderate hemisphere of PD brains (X-axis). A) entire set of 25 PD brains, B) subset of 14 least variable and clustered PD brains. Genes with FDR < 0.05 in asymmetry are plotted. C) as in Figure 1C except the comparison PD brain samples are only the 14 least variable PD brains. D) as in Figure 1D except the comparison hemispheres derive from only the 14 least variable PD brains.

Of the 9,274 significant DE asymmetry genes in the clustered subset, 61% were also DE disease genes, and of those 93% showed the opposite direction of fold-change, 7% the same direction, i.e. most genes asymmetric and higher in the severe hemisphere relative, were downregulated in PD relative to control. This is unexpected, as we predicted that the expression signature in the *hemisphere* with greater PD symptoms would correspond to the expression of the *population* with greater PD symptoms. At least in this set of most similar PD brains that relationship is reversed. Why the expression changes are inverted relative to expectation demands further discussion. However, disregarding, for now, directionality, the high correlation suggests that the same processes are affected in both comparisons. That is: genes that distinguish a severely affected hemisphere from a moderately affected hemisphere in a PD patient are largely the same as those which distinguish a healthy brain vs a PD brain, generally.

Conversely, we searched for genes that were clearly affected by (or predicted by) the side of onset but not disease status (DE asymmetry genes that were significantly asymmetric and showed at least a 0.5 log fold change (FC) *difference* in effect size between comparisons). These gene types were not plentiful but were enriched for brain that expressed x-linked (*BEX*) family genes. For the *BEX* genes identified, no difference in expression was seen for PD vs control, but these were more highly expressed in the moderate hemisphere compared to the severe hemisphere. It is therefore possible that *BEX* gene expression is associated with which hemisphere first develops PD symptoms. The resulting hypothesis is that *BEX* gene expression may be partially protective in the sense of delaying PD progression in a PD patient’s hemisphere, while having no impact on whether a person develops PD or not. These expression levels were higher in the less severe hemisphere, though due to the small number of PD brains examined, we cannot distinguish PD side onset DE from left vs right brain hemisphere differences alone. These *BEX1-5* were only identified within the small subset of PD patient brains selected due to overall RNA-seq similarity.

Some genes were more equally expressed between hemispheres than expected based on disease DE. That is, some genes that differed between healthy and disease brains appeared similar across hemispheres. Most enriched in this category were dynein genes, including 3 of 4 *ODAD* genes. Dynein related genes were DE according to disease status in both the entire comparison as well as the filtered subset.

As well as classifying asymmetry genes according to known PD disease associations, we also proceeded in the other direction: we looked for patterns of asymmetry in PD gene expression levels We considered whether variation in asymmetry of PD gene expression could corresponds to differences in disease states. The most statistically enriched PD related biological processes in this study included the genes encoding the chaperonin *CCT* (a.k.a the TriC complex). This complex of genes was significantly more expressed in PD patients vs. controls, especially in males. By generalized linear modeling we found genes correlated with *CCT* expression. These were enriched for several types of neuronal differentiation and metabolic pathways (Fig3A). This suggests that different levels of CCT could correspond to processes important in PD resistance. Indeed, in examining the distribution of *CCT* expression between hemispheres in the complete set of samples, there appeared to be a trend towards elevated expression of *CCT* genes in the less effected PD hemisphere (Supplemental Fig1F,G). This relationship was not specific to *CCT* though, and overall, there was a nonsignificant trend: the more strongly disease-associated a gene expression level was on average, the more likely it was to be expressed more in the moderately affected hemisphere. Still, these observations suggest that a pattern of asymmetry may be present in the PD genes identified.

**Figure 3.**
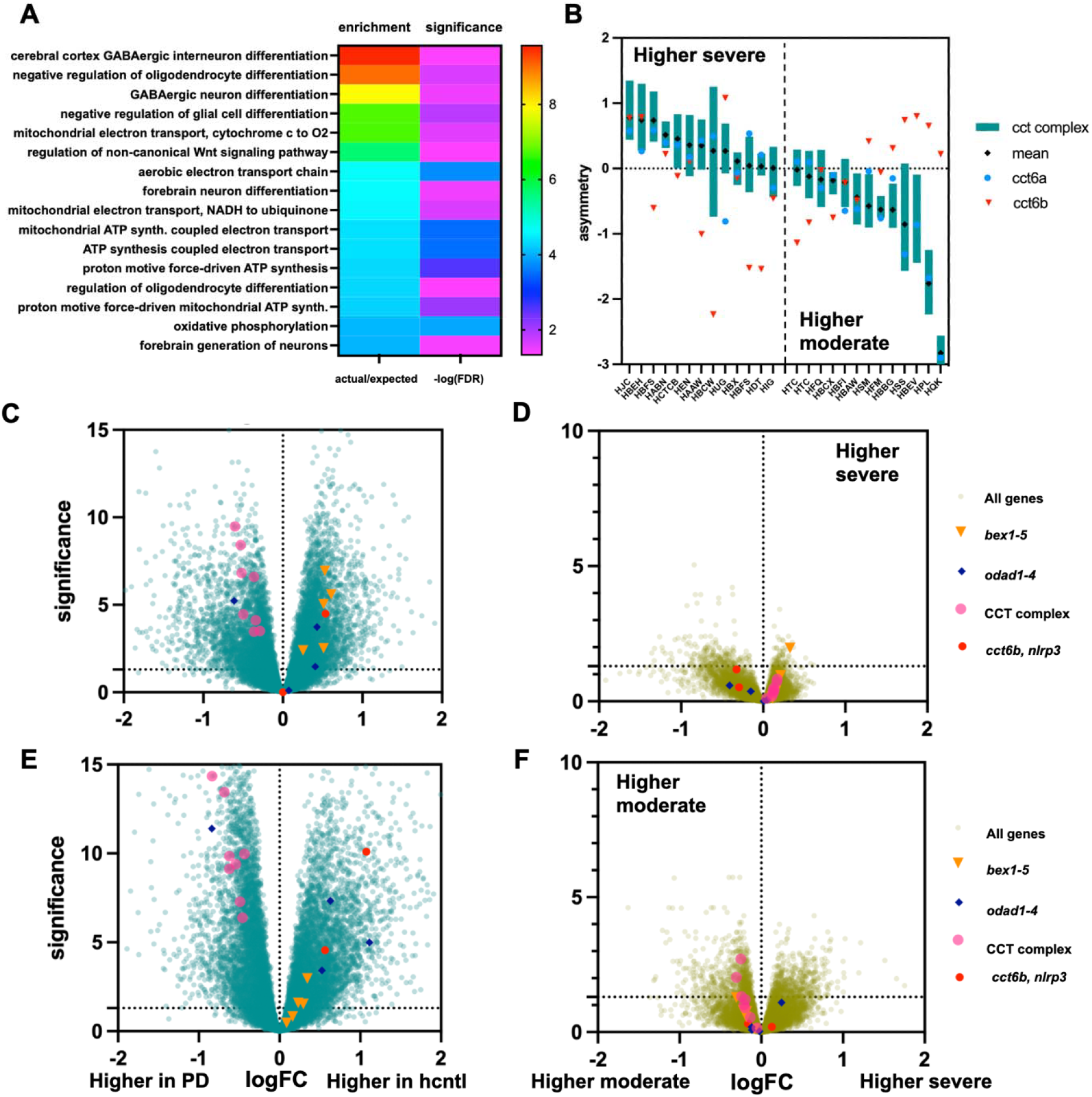
Categorizing PD brains by directional asymmetry of CCT gene expression. A) Gene set enrichment of genes with a significant association by GLM with the average expression of the 8 CCT subunit genes. B) relative expression between hemispheres of all 25 PD brains (2 samples are replicated in the plot, bars represent mean and standard deviation of the 8 CCT genes by zscore exp.severe – moderate). To the left of the vertical dashed line are the samples with higher CCT expression in the severe hemisphere, to the left those with higher in the moderate hemisphere. C) Volcano of [CCT higher in severe]-PD vs all controls. D) Volcano of severe vs. moderate expression levels in the [CCT higher in severe]-PD set. E) Volcano of [CCT higher in moderate]-PD vs all controls. D) Volcano of severe vs. moderate expression levels in the [CCT higher in moderate]-PD set.

We noticed one other interesting feature in *CCT* expression related to the alternative subunit, *CCT6b*. The significant disease associated *CCT* genes we identified include *CCT6b* which has previously been reported to be inversely correlated with the other *CCT* gene subunits, especially with respect to cancers [21, 22]. In agreement with this observation, in the present study, genes *CCT1-8* were more highly expressed in healthy control tissue whereas *CCT6b* was proportionally less expressed, relative to PD patient brains. But comparing across hemispheres within patients, *CCT6b* was on average similarly expressed to the rest of the *CCT* subunits, apparently pointing to inverse correlation by disease but not hemisphere (Fig3B). When measured explicitly though there was, in fact, a negative correlation by cpm (r^2 = -0.44) or no correlation according to normalized counts (r^2 =0.0) of *CCT6b* with the primary subunits compared to high positive correlation between the primary *CCT* genes (cpm: r^2 = 0.93, normalized: r^2 =0.85). The explanation is, of course, that whereas the average expression of the CCT6b and the primary *CCT* subunits were similar, for given PD individuals the expression profiles were often negatively correlated, but with many people showing asymmetry in one direction and other in the opposite direction (Fig3B). That is, there was a continuum to the expression of *CCT* genes, with some PD patients showing increased expression in the severe hemisphere and some showing increased *CCT* expression in the moderate hemisphere, with the alternative subunit *CCT6b* being weakly to negatively correlated correspondingly. We asked first whether statistically significant gene expression asymmetry was present after stratifying PD patients according to *CCT* gene expression asymmetry. Incidentally this method of categorization produced a different grouping than the clustering-based approach in Fig2.

After stratifying all PD patients based on the directionality of CCT gene asymmetry, similar DE was observed for both types compared to healthy controls. However, significant asymmetry in gene expression was seen for both groups, with many genes having opposite directions of asymmetry and other genes showing asymmetry in one group but not the other (Fig3C-F). Comparing the gene set enrichment of significant genes produced different ontologies and a direct comparison of the two PD sets yielded hundreds of DE genes (Supplemental Fig3, Supplemental Data 1,2). The top 25 small (>500 gene) biological process GO categories that are enriched in DE genes (and excluding those distinguishing PD from WT) include three categories that may relate to CCT activity directly, “tube morphogenesis”, “supramolecular fiber organization”, and “response to wounding.” Seventeen of the 25 are related to immune function, including “Innate immune response”, “Leukocyte activation”, “Regulation of lymphocyte activation”, and “Regulation of T cell activation”. It is possible that CCT asymmetry differences are related to different immune response in in PD.

## Discussion

General brain hemispheric asymmetry in neurodegenerative diseases was reviewed in a publication by our group [2]. It formed the basis for the present examination of differences in gene expression between hemispheres within PD patients and in comparison, to healthy control brain samples.

Three classes of genes were identified. These include: DE genes associated with both Parkinson’s disease and hemisphere relative to side of PD onset (including the *CCT* genes), DE genes associated with disease status but not hemispherical levels (including dynein genes and *ODAD1-4*), and DE genes associated with hemisphere but not disease (including *BEX1-5*). These classes support the possibility that some biological processes are distinctly asymmetrical before and/or during the etiology of the PD, causing or the result of disease.

The CCT chaperonin complex is associated with PD and can modulate disease processes [23], and is required for proper folding of tubulin, is a modulator of protein aggregation, and may impact immune processes [24, 25]. Brain expressed x-linked (BEX) genes are highly expressed in neuronal tissue, have been linked to stem cell capacity, and is upregulated in brain tumors cancers [26-29]. Both gene sets were more highly expressed in the less affected hemisphere of similar PD brains. The variation could be either protective or indicative of reduced damage by the disease.

Counter to expectation the *CCT* genes showed anti-correlation between disease status and hemisphere expression, with the less affected hemisphere having *higher CCT* expression compared to the severe hemisphere, even though healthy patients (less affected by PD) had a lower *CCT* expression compared to PD patients. This paradox could be resolved in various ways. For instance, CCT-mediated processes could be both compensatory at a local level and unregulated globally in response to PD. Thus, one hemisphere could exhibit more effective compensation and so reduced markers of PD pathology due to greater levels of CCT. Interestingly, control female brains, here, had high *CCT* expression levels while healthy male brains had low levels. In the PD brains *CCT* expression was roughly the same for male and females and more like control female levels. This suggests that CCT might be protective at a certain level with healthy females already expressing that level, but males have ‘room’ for increased expression. Suggestively, it is known that men have from 1.5 to 2-fold the rate of PD diagnosis as women [30].

We conjecture that at least 2 PD subtypes are present in the patients in this study, and these can be typed according to *CCT* expression patterns. One subtype has increased expression in the severe PD hemisphere the other increased expression in the less affected hemisphere. Both correspond to a set of differentially expressed but different genes. A speculative biological explanation does require a somewhat complicated model where the CCT chaperonin is involved in both PD types but in different ways or to different degrees. This is possible and hypothetical scenarios can be readily imagined. For instance, one PD type may be driven by alpha-synuclein misfolding, which is impeded by compensatory chaperonin presence, whereas in another increased neural cell body growth or even limited proliferation is associated with lower levels of CCT overall and increased levels of *CCT6b* expression.

Among the healthy control brains, the inverse relationship for CCT6b expression and the primary CCT subunits was also present, 4/10 vs. 12/24 for the PD case brains. However, the degree of asymmetry for these genes was greater in the PD brains. It may be that asymmetry is present but is exaggerated following disease onset. Alternatively greater asymmetry may make PD more likely. This remains highly speculative at this stage.

Gene expression differences between the severe and moderate hemispheres within PD patients were identifiable following stratification of PD patients. Most of these genes correspond to and were also identified by comparing PD brains vs. healthy controls in general. It is strange, and worth further study, as to why the direction of expression is inverted by side of onset compared to disease state. We suspect the origin of this pattern could be related to compensatory effects which are initiated in a disease state, but which are more apparent in the less effected hemisphere due to less large-scale disruption. The brain samples derived from prefrontal cortex and from relatively early-stage PD, mostly Braak stages 1-3 (a single stage 5, and 1 stage 6) [9]. We would not expect severe neurodegeneration in this region for these patients. Thus, we hypothesize that important cellular stress related processes can both permeate the entire brain and/or be partially bound to the originating hemisphere.

PD is a complex neurodegenerative disorder with only partially understood underpinnings. Ultimately the present study compared gene expression from 28 patients and 10 control brains. Meaningful comparison between hemispheres within patients required additional stratification of PD cases. Limited biographical information associated with the samples unfortunately precludes analysis or even speculation regarding what else, beyond gene expression patterns, differentiated these groups of patients. We also cannot extrapolate how representative these patients may be for PD patients overall. For these patients though, layering a comparison across hemisphere onto the greater analysis of disease required additional grouping. This could be taken to imply that PD is composed of distinct subtypes or even that the disease has intrinsically unique features for every individual. Here *CCT* genes seemed useful for discrimination. More generally, asymmetry related measurements may become useful for PD research and clinical diagnosis, but only in the context of careful filtering and calibration of disease subtypes and individual variation.

In classic twin studies, genetic, epigenetic, and environmental differences are controlled; this allows for the precise examination of the remaining non-concordant phenotypes [31]. These studies are also the gold standard for estimating the genetic heritability of traits and the relative contribution of environmental vs innate risk factors (for PD around 30%). Incidentally, these studies show slightly higher disease concordance in identical twins vs. fraternal (in one study 0.2 vs 0.13), no measurement of side of onset concordance has yet been done [32, 33]. Each hemisphere of a brain is a near identical copy of the other, differing only subtly (genetically and anatomically) due to functional lateralization and minor differences in environmental exposures. In the present study we find that side of onset is partially correlated with differences in gene expression even in regions not yet directly affected by the disease.

## Acknowledgments

The authors thank the Van Andel Flow Cytometry Core and Genomics Core for providing sorting and sequencing facilities and services. This work was supported by NIH grant R01NS113894.

