## Supplemental Figures 1-3 for "Gene expression asymmetry in Parkinson’s Disease; variation of *CCT* and *BEX* gene expression levels are correlated with hemisphere specific severity"

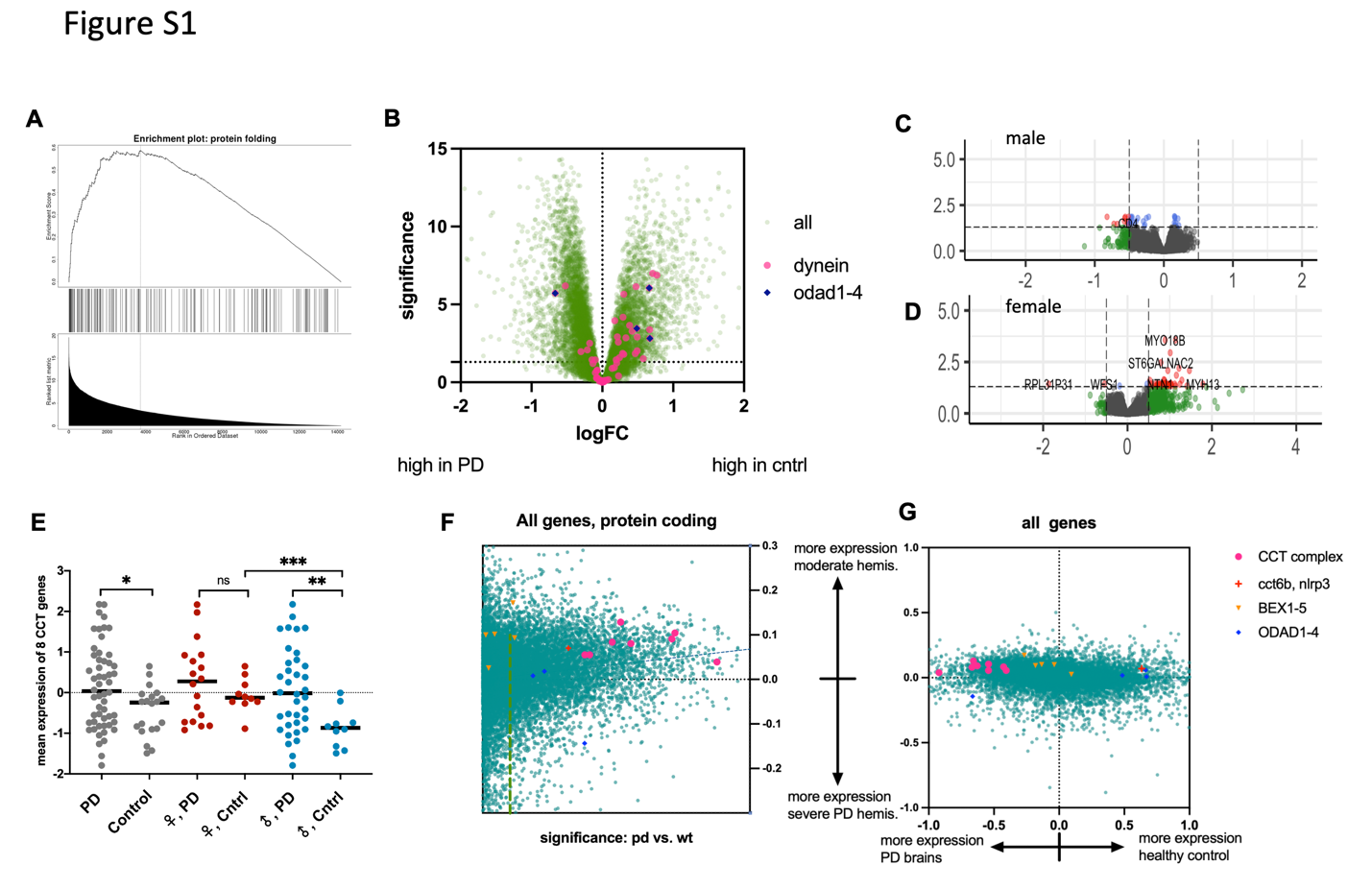

***Supplemental Figure 1. A) ranked gene set enrichment analysis, ordering is by significance comparing PD to healthy controls. B) Volcano plot of PD vs healthy controls with all genes named or related to dynein indicated in magenta. C) Volcano plot of male PD brains vs. male healthy controls. D) Volcano plot of female PD brains vs. female healthy controls. E) comparison of CCT gene expression. Expression is in Z-score normalized RUV-seq adjusted counts. F) asymmetry logFC (y-axis) vs significance of PD vs healthy control. G) asymmetry logFC (y-axis) vs. logFC difference healthy vs. PD.***

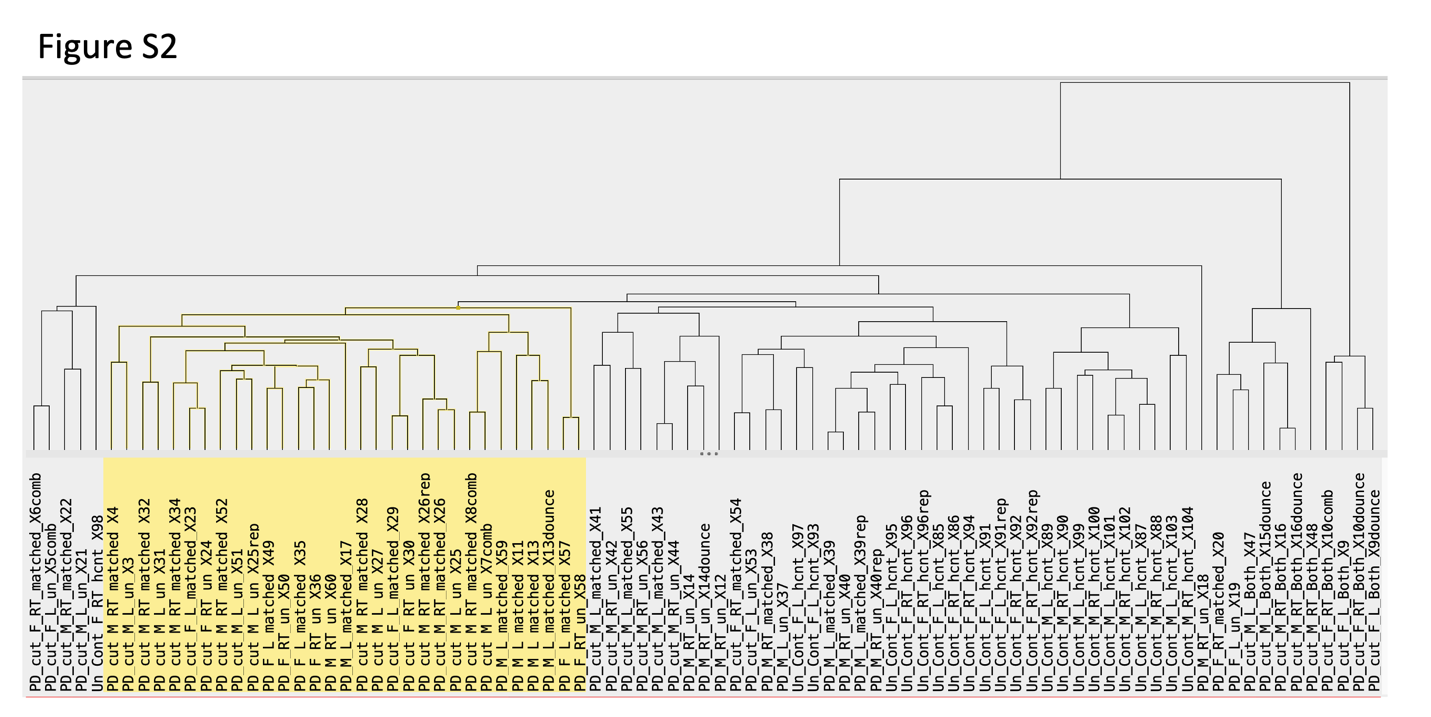

***Supplemental Figure 2. Clustering (by average) of all samples by rank correlation of the top 2,500 most variable high expression genes. The most similar 14 PD patients (8 males, 6 females, indicated in yellow) were compared to 10 control brain samples.***

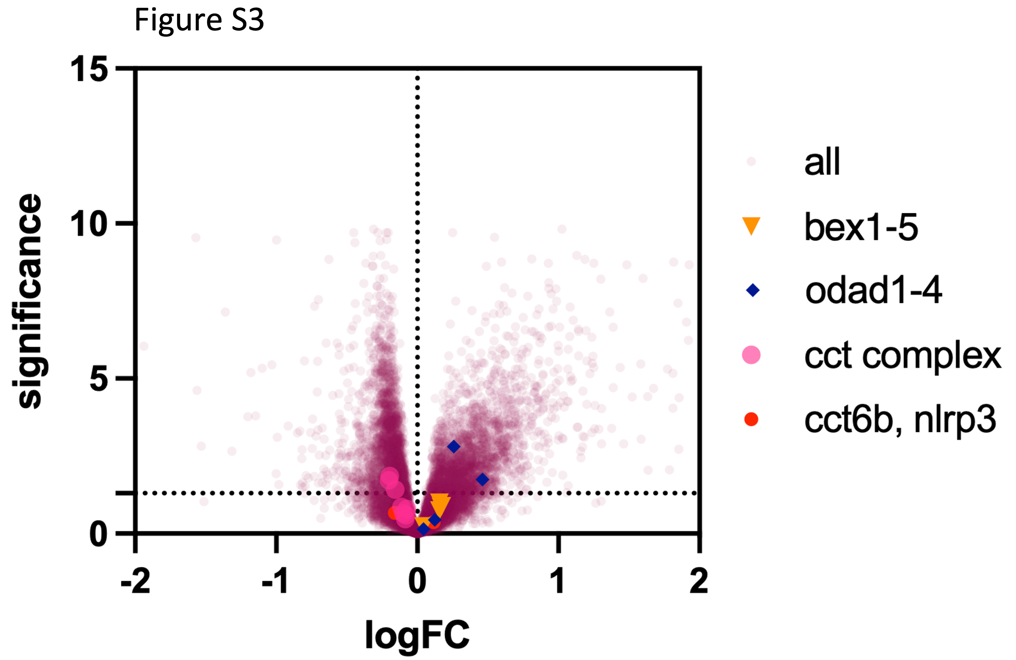

***Supplemental Figure 3. Volcano plot comparing PD patients with asymmetrically higher* CCT *expression in the severe hemisphere to those with higher* CCT *expression in the moderate hemisphere.***
